# Integrating natural language processing and genome analysis enables accurate bacterial phenotype prediction

**DOI:** 10.1101/2024.12.07.627346

**Authors:** Daniel Gómez-Pérez, Alexander Keller

## Abstract

Understanding microbial phenotypes from genomic data is crucial in areas of research including co-evolution, ecology and pathology. This study proposes a new approach to integrate literature-derived information with genomic data to study microbial traits, combining natural language processing (NLP) with functional genome analysis. We applied this methodology to publicly available data to overcome current limitations and provide novel insights into microbial phenotype prediction.

We fine-tuned specialized transformer-based large language models to analyze 3.3 million open-access scientific articles, extracting a network of phenotypic information linked to bacterial strains. The network maps relationships between bacterial strains and traits such as pathogenicity, metabolic capacity, and host and biome preference. By functionally annotating reference genome assemblies for strains in the phenotypic network, we were able to predict key genes influencing phenotypes.

Our findings align with known phenotypes and reveal novel correlations, leading to the identification of microbial genes relevant in particular disease and host-association phenotypes. The interconnectivity of strains within the network provided further understanding of microbial community interactions, leading to the identification of hub species by inferring trophic connections—insights challenging to extract by means of experimental work.

This study demonstrates the potential of machine learning methods to uncover cross-species patterns in microbial gene-phenotype correlations. As the number of sequenced strains and literature descriptions grows exponentially, such methods become crucial for extracting meaningful information and advancing microbiology research.

## 1. Introduction

The quest to decode the intricate relationship between an organism’s genome and its phenotype has been a cornerstone of biological research since the discovery of DNA as the source of heritable material (Elison & Acar, 2018; Rahaman et al., 2015; Wong et al., 2021). Contrary to initial expectations, the link between phenotype and genotype remains complex and challenging to unravel as there is not a one-to-one relationship between phenotypes and genes (Fisch, 2017). Rather, most phenotypes are polygenic, depending on multiple genes for expression (Boyle et al., 2017; Visscher et al., 2017). Adding to this complexity, the genomic context of a gene and epigenetic factors also play a role in the expression of distinct phenotypes from the same genomic material (Atkinson & Halfon, 2014; Feinberg & Fallin, 2015). Despite the increasing wealth of publicly available genomic data, the challenge of predicting accurate microbial phenotypes from the relatively small and low-complexity bacterial genomes persists, presenting a critical bottleneck in fields ranging from epidemiology to environmental science. This study introduces a novel approach to this enduring problem, leveraging the potential of natural language processing (NLP) and machine learning.

Despite hundreds of thousands of high-quality sequenced prokaryotic genomes being available in public databases such as NCBI, a significant challenge remains in the form of limited accompanying metadata (Haft et al., 2023). The latter, which includes information such as environmental and growth conditions, as well as phenotypic characteristics, is crucial for a comprehensive understanding of the genomic data. As a result, genetic data, while extensive, cannot be fully leveraged or properly interpreted without this contextual information. This limitation restricts the potential for meaningful insights in areas such as host interaction, metabolic assessment, and environmental adaptability of bacteria. Although databases exist that tackle this problem, they are mostly focused on metabolite usage and lack more complex phenotype annotations (Dérozier et al., 2023; Söhngen et al., 2014). To address this gap, there is an urgent need for more systematic and thorough documentation of phenotypic traits alongside genomic sequences as well as more efficient tools to synthesize and summarize them (Keller et al., 2023). Such advancements would significantly enhance our ability to harness the full potential of genomic datasets, leading to more accurate predictions of bacterial behavior and responses under varying conditions.

Recent progress in NLP, including the development and high popularity of large language models (LLMs) based on transformer architecture, has opened new avenues in biological research (Madani et al., 2023; Singhal et al., 2023). Specifically, models based on the BERT architecture pre-trained on scientific texts such as SciBERT (Beltagy et al., 2019) and BioBERT (Lee et al., 2019) have been fine-tuned to perform tasks such as named entity recognition (NER) and relation extraction (RE) to obtain meaningful information from literature. Compared to foundational models such as ChatGPT, these specific models have been shown to perform better in benchmarks for data mining tasks (Karkera et al., 2023). More recently, BioLinkBERT (Yasunaga et al., 2022), a BERT-type model trained on hyperlinks connecting scientific articles, has shown the best overall performance on the Biomedical Language Understanding & Reasoning Benchmark, BLURB (Gu et al., 2021). BLURB includes datasets for NER and RE tasks such as NCBI-disease (Dŏgan et al., 2014) and Drug-Drug Interactions (Herrero-Zazo et al., 2013), respectively. These are straightforward problems with a small number of different types of entities and relations that serve as benchmark and proof-of-concept. However, complex multimodal implementations leading to novel insights are lacking in the literature. In particular, the problem of phenotype-genotype correlations has not yet been tackled using such approaches outside the medical domain. Encoder-only transformer models such as BioLinkBERT offer a promising solution to the long-standing challenge of linking genotypic information with phenotypic traits by easing literature mining and interpretation.

This work aims to bridge the gap between genomic datasets and limited phenotypic metadata. We leverage the power of NLP, specifically fine-tuned BioLinkBERT models, to systematically extract information from the vast literary corpus of PubMed Central (PMC) and paint a picture of the microbial phenotype landscape in scientific literature. By integrating this extracted knowledge with functional genome annotations from public databases, we investigate correlations between bacterial phenotypes and their underlying genetic makeup. We additionally show several examples of using this network to extract biological insights, including the inference of microbe-microbe interactions and gene-host correlations. This study expands the applications of cutting-edge NLP techniques in biology, ultimately leading to new insights and a deeper understanding of microbial life.

## 2. Methods

### 2.a. Data and model preparation

#### 2.a.i. Database

The PMC database was selected as the source for the literature corpus due to its comprehensive and up-to-date collection of open-access scientific articles together with license information for data mining. Article data were retrieved from the PMC release date 2024-06-18, including all commercial and non-commercial open-access datasets. After filtering out journals not specifically related to biology and microbiology, sentence tokenization (breaking text into sentences) and section classification were performed using the R package tinypmc. Sentences shorter than 40 or longer than 512 characters were discarded, along with those from sections unlikely to contain strain descriptions, including acknowledgements, author declarations, data availability statements and legends. The sentence length cutoffs were designed for optimized speed while minimizing the loss of phenotypic information, based on preliminary tests. Additionally, duplicated sentences and those in languages other than English were discarded based on predictions from the lingua-py package (confidence in English language <40%). We replaced the hyphen in all hyphenated words with a space to avoid token splitting in the NER models.

#### 2.a.ii. Annotation

For the annotation task, we defined entity groups to extract microbial phenotype information. They belonged to the following categories for classification: taxonomic (STRAIN, SPECIES, ORGANISM), phenotypic (PHENOTYPE, EFFECT, DISEASE), and environment/molecule-related entities (MEDIUM, ISOLATE, COMPOUND). STRAIN and SPECIES exclusively included properly characterized bacterial strains and species. All other taxonomic classifications, including fungal strains or animal species, were grouped under ORGANISM. The phenotypic entities were classified into PHENOTYPE, including for example the shape of bacteria or trophic characteristics. EFFECT contained a variety of entity classes indirectly related to effects caused by strains that did not fit into the PHENOTYPE category. DISEASE contained the names of diseases or pathological signs. COMPOUND included organic and inorganic molecules, such as common metabolites, metals and antibiotics. Finally, MEDIUM was related to the growth medium of bacteria in laboratory settings, and ISOLATE to the type of sample from which the particular bacterial strain was acquired.

Regarding relation categories to connect the previous entities, we defined the following: interaction with environment (GROWS_ON, INHABITS), metabolic interactions (PRODUCES, DEGRADES, RESISTS), biological phenotypes (PRESENTS, ASSOCIATED_WITH, PROMOTES) and interactions with other organisms (SYMBIONT_OF, INFECTS, INHIBITS). SYMBIONT_OF group, being a subcategory of INHABITS, had both labels. The links were not restricted to particular entities but followed logical sense, resulting in cases where the same relation connected different kinds of entities (i.e., INHIBITS, was both used to connect STRAIN and ORGANISM as well as COMPOUND and STRAIN). However, in some cases the entity was connected through an exclusive relation type (i.e., GROWS_ON, for MEDIUM). For clarity, we refer to specific RE categories in this document by specifying first the two entities linked by the RE in the direction of the connection with a dash, followed by a colon and the RE category name (i.e., sourceNER-targetNER:RE).

Initially, we performed a manual annotation of entities and their relations on a set of randomly selected sentences that contained popular strains matched by their exact strain name as found in the literature corpus. With these initial annotations, we trained preliminary models and ran them on a small subset of the full corpus. In order to improve the models, we iteratively selected and annotated new sentences containing predicted labels in cases where they differed widely from the ground truth. In total, we annotated 3,798 sentences. The annotated dataset included sentences with both presence or absence of each of the entities. Since the base rate for the presence of strains in random sentences from the literature is relatively low, a large number of non-entity-containing sentences (≈35.1%) were included to prevent false positives. Common variations of some of the most abundant entities after NER prediction were added by manual curation to improve downstream analyses.

#### 2.a.iii. Model fine-tuning

We fine-tuned a base BioLinkBERT-large model (24 encoder layers, 16 attention heads, 340M parameters) for each of the categories with the task of entity recognition, resulting in 9 separate single-NER prediction models. A stratified split (same proportion of labels on each set) based on all NER annotations was performed, resulting in 65%, 17.5% and 17.5% of the total sentences, corresponding to training, validation and testing, respectively. For each entity, a Beginning-Inside-Outside (BIO) encoding was applied to the training sentences and each of the 9 models was fine-tuned for at least 20 epochs with early stopping of 5 epochs at a learning rate of 2 × 10^−^5. BIO encoding consists of three label classes *C* which takes class *c* for each token *t* in a sentence (*c* ∈ {*B*, *I*, *O*}). If the token is at start of the entity, its class is *B* and each continuing token is *I*, all others outside of the entity are labeled as *O*. The best models were selected based on the minimization of cross-entropy loss *L* in the evaluation set for each entity. For the set *N* of sentences in the training dataset, the loss was calculated as:

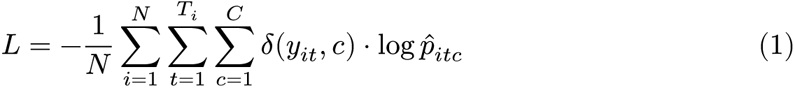

where *T*_*i*_ is the number of tokens in the *i*-th sentence and *y*_*it*_ is the true label for the *t*-th token in the *i*-th sentence. The *δ*(*y*_*it*_, *c*) corresponds to Kronecker delta function, and is 1 if the label *y*_*it*_ equals class *c*, and 0 otherwise. The AdamW optimizer was employed for gradient descent as implemented in the Hugging Face transformers package (Loshchilov & Hutter, 2017).

Regarding RE models, we prepared a dataset for each relation type consisting of sentences annotated with binary labels indicating the presence or absence of the relation between the specified entities. We split all pairwise relations into 60% training, 20% evaluation and 20% testing, performing stratification in this case per individual relation, due to disparities in entity distributions of the dataset. For the training of RE models, we used a similar strategy as for the NER models, but optimizing instead for a higher F1 score in the evaluation dataset, the harmonic mean of precision (*P*) and recall (*R*):

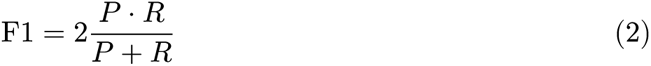

for a total of at least 15 training epochs with early stopping and a learning rate of 3 × 10^−^5. All training was done on four Nvidia A100 Tensor Core GPUs (80GB memory) in parallel using the PyTorch backend (Paszke et al., 2019).

For the prediction pipeline, we initially used the STRAIN model to annotate all relevant sentences from the filtered corpus in the PMC database. In the NER annotation, we considered a positive hit when the prediction score was 0.5 or higher (*p*(*t*_*i*_, *c*) ≥ 0.5) after aggregation of consecutive tokens (*T*_*c*_ = (*t*_*k*_, *t*_*k*+1_…*t*_*n*_)). Token aggregation was performed using the max aggregation score from the Hugging Face transformers package which results in the entity assignment with the maximum score for a series of *T*_*c*_ tokens when *c* ∈ {*B*, *I*}:

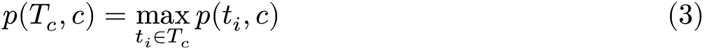

(Wolf et al., 2019). On the resulting positive sentences for the presence of a STRAIN entity, we performed predictions with the remaining 8 NER models. Finally, we applied the RE models for extracting relations on sentences with at least two NERs, one of which corresponding to a strain, by pairing all combinations of STRAIN with other NER categories within the same sentence. We chose those with a score over 0.5 for a positive RE label and compiled the annotations. The threshold of 0.5 in the NER and RE classification was chosen due to standard practice in machine learning, and led to a good balance of false positive minimization. Due to the similar prevalence of classes in the training and corpus, the cutoff was left in place.

For visualizing the learned representations of the models, we used the UMAP dimensionality reduction algorithm (McInnes et al., 2018). For sentences with either presence of at least a positive entity in the NER models or a positive relation with RE, we applied UMAP on the per-sentence mean token outputs from the last hidden layer, resulting in a 1024-dimensional vector for each sentence. UMAP parameters used were as default except for the minimum distance being 0.01. This allowed better visualization of the local differences as opposed to the global picture, which was more informative in this case.

### 2.b. Network analysis

#### 2.b.i. Network construction

To analyze the interconnectivity between entities and relations, we represented the extracted information as a directed network *G* = (*V*, *E*), where *V* is the set of nodes representing entities, and *E* is the set of edges representing the relations between entities. The degree distribution *P* (*k*) of the network, where *k* is the degree of a node (the number of connections it has), was assessed to determine if the network exhibits a scale-free property, which follows a power-law distribution:

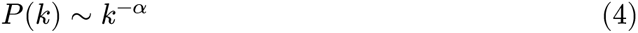

where *α* is the scaling exponent. Algorithms for the network analyses, including distance metrics and triad assessment, were applied using the Graph-tool and Networkx packages in Python (Hagberg et al., 2008).

#### 2.b.ii. Community interaction

From the subnetwork constructed of strains and compounds associated with the same environment, we considered two types of negative strain correlations. Either when two different strains degraded the same compound or when a compound produced by one strain was responsible for an INHIBITS relation towards another strain, as an example of resource competition and direct inhibition, respectively. Positive correlations between strains were defined by the degradation of a product by a strain that is produced by another to exemplify cross-feeding, as well as RESISTS relation of a strain towards a product produced by another strain to represent a resistance-development-type interaction. For each pair of strains in the subnetwork (*s*_*i*_, *s*_*j*_) we calculated the interaction weight *w*_*ij*_:

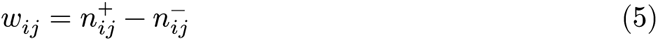

where 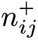 and 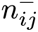 is the number of positive and negative interactions inferred between strains *s*_*i*_ and *s*_*j*_, respectively. The interaction weight *w*_*ij*_ provides a quantitative measure of the overall relationship between two strains, with positive values indicating net positive interactions and negative values indicating net negative interactions.

### 2.c. Phenotype-genotype correlation analysis

To handle potential biases and disambiguations in the name of the strains, we matched the resulting predicted values in the STRAIN label to the Strainselect reference database, which links different nomenclatures for the same strain, using a custom approach based on Levenshtein string distances (DeSantis et al., 2023; Wagner & Fischer, 1974). This included first a local distance match to assess partial strain names due to the common presence of abbreviations, followed by a full distance assessment to map complete names. We performed a similar approach on non-STRAIN entities in order to group variations of the same entity, such as plural forms or typos. The representative entity name was chosen based on the largest number of occurrences in the literature.

We downloaded the NCBI RefSeq strains connected to each of the strain matches. For each assembly, we predicted the proteome using the Prodigal prokaryotic annotation tool (Camacho et al., 2009). We performed *de novo* functional annotation using Interproscan 5.67 based on homology to Pfam families. (Jones et al., 2014; Mistry et al., 2020). We then encoded all functional annotations using their abundance (number of copies for an annotation) in that particular assembly as features and grouped all strains that had the same phenotype relation in a matrix.

Enrichment of GO terms was calculated using Fisher’s exact test and correcting p values for multiple testing using false discovery rate as implemented in GOATOOLS (Klopfenstein et al., 2018). GO release 2024-06-17 was employed. Species phylogenies were inferred using the Open Tree of Life API Python package (Mctavish et al., 2021).

#### 2.c.i. Gradient boosting

After discarding single-appearance entity terms, we performed a stratified split to separate the train and test sets (70% and 30%, respectively) for each of the 17 unique relations between entities. We used a gradient boosting approach for binary classification of each of the labels that were found in 10 or more assemblies from two or more different genera with early stopping using the XGBoost library (Krishnapuram et al., 2016). For models with accuracy higher than 80%, we extracted the importance gain, corresponding to the average improvement in the training set loss brought by a particular feature. We used this to rank how relevant the functional features were for the corresponding phenotype labels and correlations.

#### 2.c.ii. Positive selection analyses

To study the selective pressures of high-importance genes, we selected gene annotations belonging to strains within a specific relation category that had the same annotation. We aligned these using MAFFT and assessed presence and significance of gene-level positive selection for annotations of each relation category using BUSTED (Katoh & Standley, 2013; Murrell et al., 2015). BUSTED is based on the ratio of nonsynonymous (*d*_*N*_) to synonymous (*d*_*S*_) amino acid change to infer selective pressure:

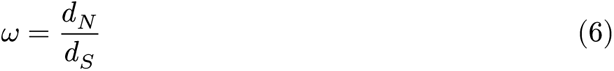

where *ω* < 1 is purifying selection, and *ω* > 1 indicates positive selection. BUSTED fits each gene to a constrained null model (*M*_0_) that does not allow for positive selection (*ω* ≤ 1) and an alternative unconstrained model (*M*_1_) that allows for positive selection (*ω* > 1). Likelihood ratio test (LRT) statistic is:

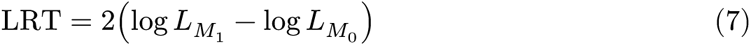

where 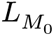 and 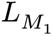 are the likelihoods of each model, respectively. A significant LRT value suggests evidence for positive selection.

### 3. Results and discussion

### 3.a. Phenotype data extraction from literature

#### 3.a.i. Annotations

We extracted microbial phenotype information from the literature corpus using the bioinformatics workflow described in Figure 1a. The entity annotation in the training dataset was meant to represent the frequency of strain-phenotype information in the literature since the approach at the beginning of the manual annotation was to randomly select sentences containing exact strain matches. However, this shifted as we added more sentences to the annotated dataset to improve the models. Out of the 3,798 manually annotated sentences, 2,464 had at least one strain (≈64.9%) while the total number of individual bacterial strains annotated was about double this figure, with a similar pattern for other entities (Figure 1b). As expected, most sentences had a single STRAIN, as this was the key entity from which we aimed to extract relational information. The lowest number of entity annotations corresponded to DISEASE, which had ≈350. However, the number of entity annotations in the annotated dataset was not correlated with the number of relations to STRAIN, but rather related to how phenotype information is described in the literature. For example, the number of relations to STRAIN was larger for ISOLATE (the name of the strain is frequently reported together with where the sample was taken) than to SPECIES, which frequently is reported in a list with other strains, to which no direct relations can be drawn.

**Figure 1:**
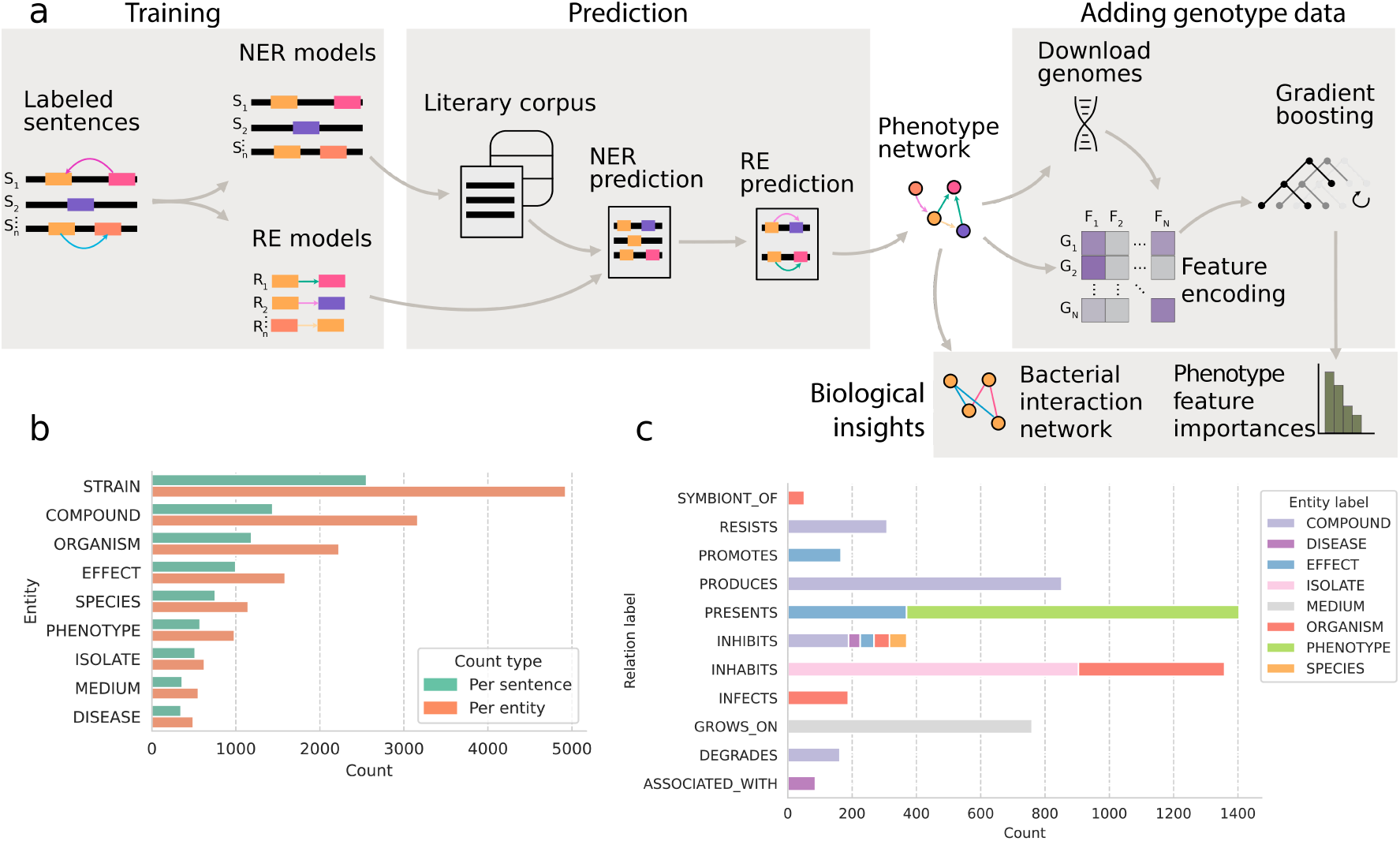
Workflow and training dataset. **a** Schematic of the bioinformatics pipeline used in this study. **b** Bar chart showing the number of entities annotated either per sentence or over the total dataset. **c** Stacked bar chart showing the total number of entity annotations per relation type. The colored bars represent different entities and their size the number of their respective occurrence in the same sentence with a STRAIN entity in the training dataset.

Relation types with the higher number of annotations included INHABITS and PRESENTS (*n* ≈1300), followed by PRODUCES and GROWS_ON with ≈800 (Figure 1c). The remainder had between 50 (ASSOCIATED_WITH) and 350 (INHIBITS) annotated sentences. Most abundant same sentence co-occurring entities with STRAIN included COMPOUND, SPECIES and ORGANISM and least abundant corresponded to MEDIUM and DISEASE, following the same pattern of entity abundances. However, more abundant entities did not necessarily have proportionally more relations to strain. The more abundant corresponded instead to MEDIUM and PHENOTYPE, while SPECIES had the least (Supp. Figure 2). As mentioned earlier, this relates to patterns of reporting strain information in scientific papers.

#### 3.a.ii. Accurate prediction across entity types

The NER models resulted in an accurate prediction of over 60% for most entities in the test (not seen before by the model) and evaluation (used to optimize weights during training) datasets, with STRAIN scoring the highest across all metrics (Figure 2a). The lower performance for EFFECT (F1 = 0.22) and ISOLATE (F1 = 0.32) may be attributed to the subjective nature of these categories. For the former, these included phenotype descriptions that did not fit classical microbial phenotypes such as indirect effects on a host. For the latter, these included very specific isolation location descriptions. The number of samples in the training dataset for both these categories may have been too small to capture this diversity and thus is reflected in the higher loss after training. Overall, the results corresponded to an accurate prediction across all entities, with accuracies comparable to other applications of fine-tuned BioLinkBERT (Yasunaga et al., 2022). Regarding RE models, we found high evaluation metrics overall for successful classification (Figure 2b). The F1 score for all 17 models was between 0.6 and 0.9 in all cases in both evaluation and test datasets, except in STRAIN-SPECIES:INHIBITS. The higher performance compared to NER prediction may be due to the type of binary classification per sentence, instead of per token. Additionally, the straightforward nature of most connections containing both a STRAIN and other entities resulted in the presence of a relation in more than half the occurrences of STRAIN with other entities (Supp. Figures 1 and 2). In general, relations with lower abundance in the annotation set correlated to lower metrics in the test set including two INHIBITS relations, but not in the evaluation set (Figure 2b). This implies that a larger manual annotation set could lead to more accurate predictions on novel data.

**Figure 2:**
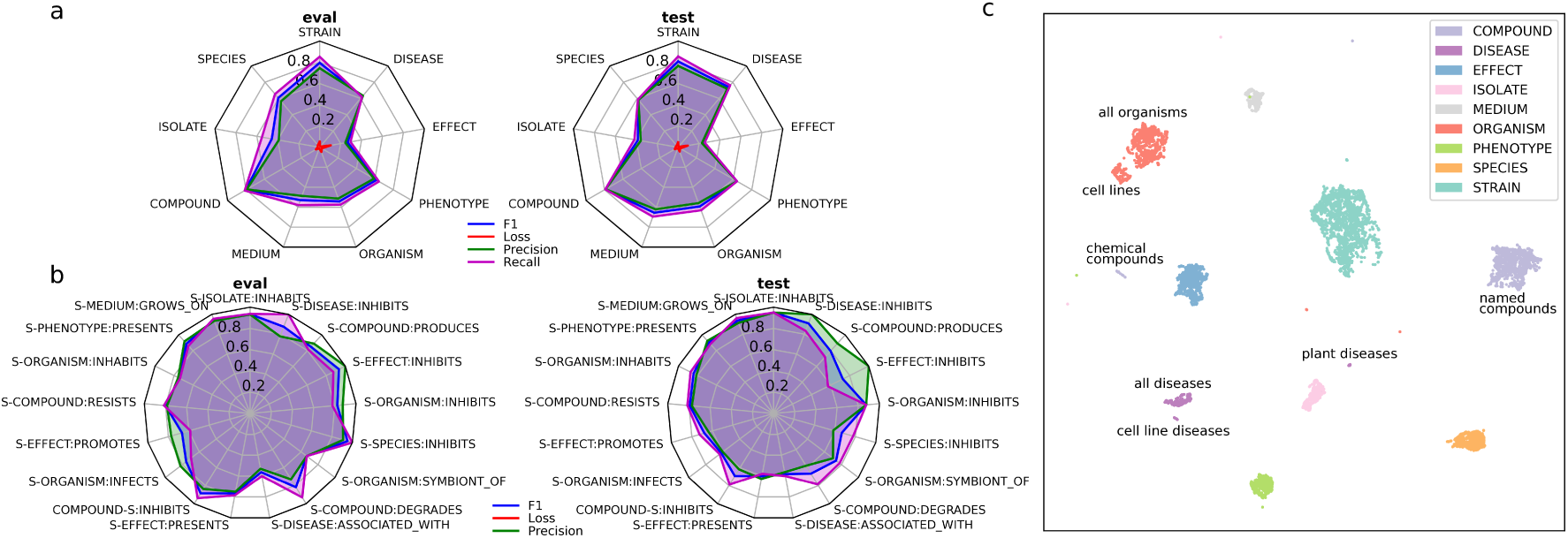
Model performance. **a** Overview of named entity recognition (NER) model training and testing. Performance metrics for the NER models categories for either the evaluation (eval) or the testing (test) data sets. **b** Overview of the relation extraction (RE) model training and testing. Performance metrics for the RE models for either the evaluation (eval) or the testing (test) data sets. S: STRAIN. **c** UMAP representation of sentence output embeddings in the annotated dataset for each of the different named entity recognition models. Displayed are only sentences that are positive for the presence of that particular entity label. Axes correspond to UMAP dimensions 1 and 2.

The lower accuracies of some NER models likely corresponded to the high variation in the number of annotated entities for training (Figure 1b), rather than misclassification of the model. Not all entity classes with low number of annotations correlated with lower prediction (Supp. Figure 3). This variability may also be related to the complexity of the particular entity and whether its complexity is captured by the examples in the annotation dataset, which is reflected in the higher cross-entropy loss after training. However, the performance difference of the NER models in the evaluation datasets is unlikely to have a large impact on the final results. Mispredictions at this stage may be filtered out by the RE models, which had overall better metrics, and the entity string matching downstream, which selected only strains with a database match, although it may have reduced the number of final correlations.

#### 3.a.iii. UMAP uncovers model representations

To understand how the models predicted the different entities, we visualized the embeddings of the 9 different NER models for the sentences from the annotated dataset in the same shared latent space (Figure 2c). In general, the embeddings showed a good separation for the different models, indicating no major overlap of the entity classes. However, for similar entities there were some subclusters that served to understand how the models see the different classes. For example, ORGANISM clusters into two, one large group containing *bona fide* organisms, and a smaller one containing cell lines, which were annotated as organisms in the dataset. Similarly, the category DISEASE formed a subcluster representing cell lines extracted from specific diseased tissues. Another small cluster for DISEASE is composed of exclusively plant diseases, and the larger one contains diseases of human and animal origin. For COMPOUND, there were also two clusters, one where the compounds are described in formulaic notation, and another larger one where they are described by the name. The clear clustering into meaningful groups was an indication of the successful grasping of nuances in the entities by the NER models.

We also visualized the embeddings of the RE models on the annotated dataset using UMAP plots. In the representations grouped by the entity (Supp. Figure 5), there is a clustering for multi-relation entities (COMPOUND, ORGANISM and EFFECT). In this case as well, there are subclusters for some types of relation. For example, in ORGANISM, there is a close association of some of the sentence embeddings from INHABITS with SYMBIONT_OF and INHIBITS. As SYMBIONT_OF and INHABITS are overlapping, this was expected, however the INHIBITS relation was surprising and may refer to the unclear distinction in many cases between positive and negative associations. Supporting this, there was an overlap of some INHABITS and INFECTS relations, which may refer to ambiguous descriptions in the literature. For the same entity there is a more heterogeneous clustering in the RE models as compared to the NER, which suggests the model learns different sentence structures that result in the same prediction. When grouping by the type of relation, the clustering is similar (Supp. Figure 4). In this case, there is a small subcluster in STRAIN-PHENOTYPE:PRESENTS which contains exclusively descriptions of bacterial colony morphology. In the INHIBITS UMAP plot, it is interesting to see how some sentences from STRAIN-EFFECT that refer in particular to antimicrobial effects cluster with STRAIN-ORGANISM. Overall, this shows the ability of fine-tuned RE models to understand different relations between entities.

#### 3.a.iv. Predictions reveal phenotype characterization imbalance

From the total ≈7,500 journals considered, 3.33 million articles were processed by the pipeline. After filtering and quality control, this resulted in a total of 510 million sentences that were considered in downstream analyses for the STRAIN prediction. Of these, 29.4 million sentences were predicted to contain at least a strain in the corpus (≈5.8%), which were fed onto the other NER models for annotation. Number of predictions for the other NERs differed slightly from the number of initial annotations. For example, the SPECIES label shifted from 3rd most abundant to 4th (Supp. Figures 1 and 2). The largest change corresponded to MEDIUM, which climbed two places to reach 5th in the prediction. Exceptions included COMPOUND (2nd most abundant in both cases) and DISEASE (least abundant in both cases). This recapitulates the baseline abundance of entities in literature, and reflects biases in the annotation, which resulted in more specific entity annotations in the training dataset due to ease of labeling and close association with STRAIN entities.

### 3.b. Phenotype information discovery using networks

#### 3.b.i. Interconnectivity of network

The annotation and prediction pipelines revealed a rich landscape of microbial phenotype information within the PMC corpus. To study the interconnectivity between all NERs and REs, we represented the information as a network where the nodes corresponded to each of the NER label categories (belonging to either STRAIN or the other 8 entities) and the directed edges to the RE correlations. The result was a large and highly interconnected network where the main component comprised most of the total nodes (≈218,000 vs. the total ≈229,100 [95.1%]) and edges (≈518,000 vs ≈525,000 [98.6%]), suggesting high connectivity between the different entities and relations.

The number of incoming and outgoing connections for each node, also known as indegree and outdegree, respectively, showed a characteristic distribution corresponding to a power-law pattern (Figure 3a). This pattern refers to the logarithmic decrease in degree with frequency of nodes. This means that while most nodes have a single connection, a minority accumulate the greatest number of edges, resulting in the formation of hubs. When fitting to a power-law, indegree has an scaling exponent *α* of 2.1 and outdegree 2.5, which indicates the more even spread of outdegree in the network. Hubs are characterized by a central node with a high degree, connected to many nodes with only a single connection. Most of these well-connected hubs corresponded to non-STRAIN entities. However, hubs where the central high-degree node is a strain are also found, and correspond to highly characterized strains in the literature. Meanwhile, the largest group of low degree nodes corresponded to strains or entities that are mentioned only once in the literature.

**Figure 3:**
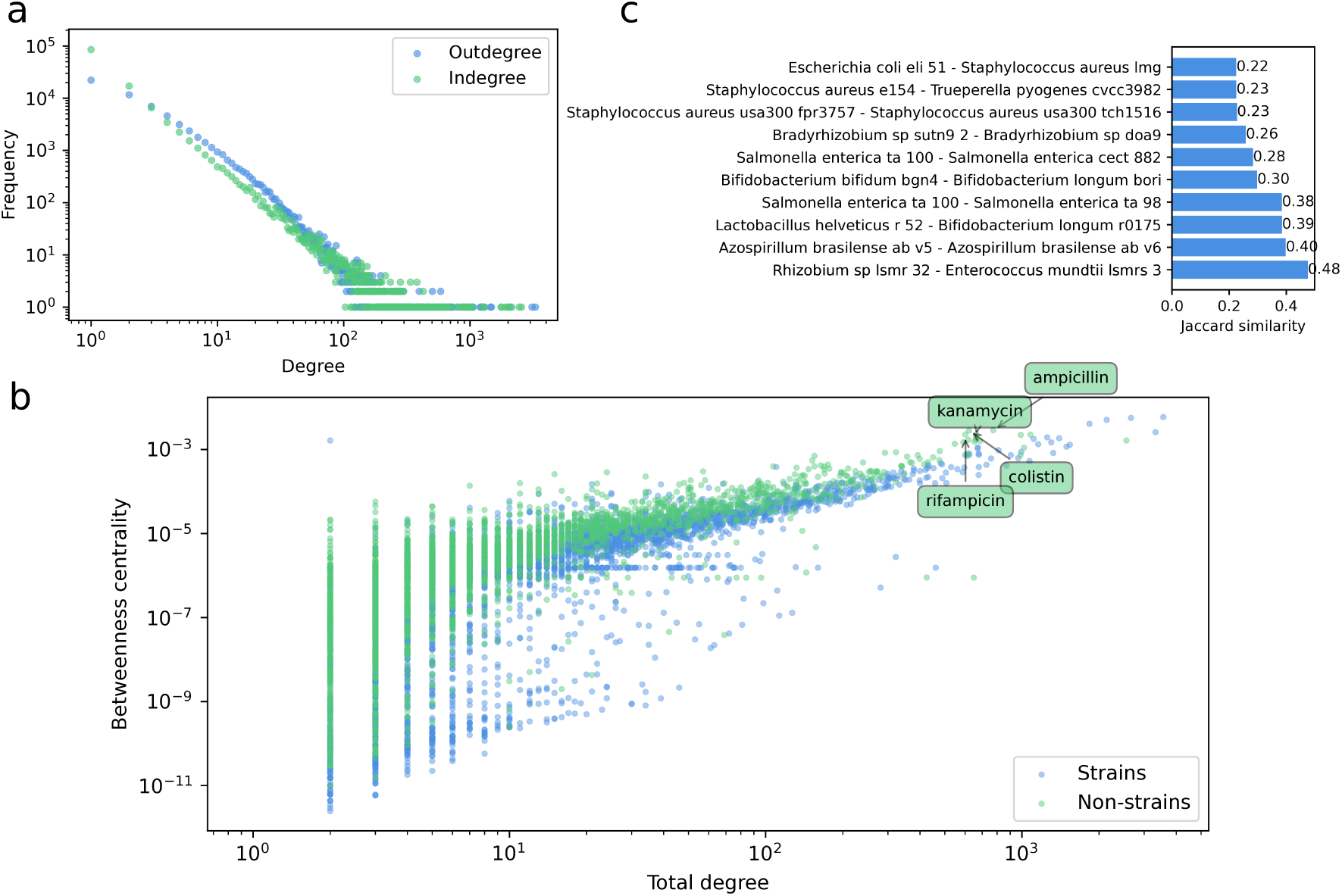
Description of phenotypic network derived from trait annotations in the literature. **a** Degree distribution of correlations in the directed phenotype network. In the x axis, the degree is represented, and in the y axis the frequency of these nodes in the network. Note the log scale on both axes. **b** Betweenness centrality of each node against its total degree. In blue are nodes of type strain and in green other node types. Labeled are the non-STRAIN nodes found in the top 10 by betweenness centrality. **c** Top 10 vertex Jaccard similarity pairs in the network.

Power-law distribution of degree occurs in graphs known as scale-free networks. These are often found in non-random biological phenomena, such as protein, gene and metabolic networks (Arita, 2005). In this particular case, the structure is likely related to the popularity and influence of certain model strains and pathogens, which accrue the most attention and publications from researchers. This, in turn, results in the hub-pattern exhibited in the network. Similarly, certain properties that are often mentioned when describing strains such as the Gram stain create hub-patterns with central nodes corresponding to these entities. The scale-free pattern was also evident in the community structure, which, when measured with the Louvain/Leiden score, resulted in a high modularity (≈0.61), indicating more links within communities than among them. This hub pattern highlights the need to develop methods to extract information which capitalize on this imbalance in phenotype descriptions. Given this structure, future work may consider using phylogenetic relationships to highlight taxonomy-specific correlations.

Nodes with the highest outdegree included only bacterial strains (Table 1), mostly *Escherichia coli*, as it is one of the most popular microbes in research (Blount, 2015). However, others relevant in, for example, pathogenicity (*Pseudomonas aeruginosa* PAO1, a model pathogen) were also highly ranked. The indegree ranking was more varied in regards to entity category, owing to the single relation type where STRAIN is acting as the target node (COMPOUND-STRAIN:INHIBITS; Table 2). Gram stain was the 3rd and 5th highest ranked term as one of the most common phenotype characteristics reported when describing bacterial strains in literature. When examining the betweenness centrality, the top nodes are again some of the most popular strains together with antibiotics, including ampicillin, kanamycin, colistin and rifampicin (Figure 3b). This suggests that these antibiotics, due to their broad spectrum of activity against different strains, may act as the main bridges linking different high-degree hubs together within the network.

**Table 1:** Top 10 nodes by outdegree in the phenotypic network.

| Node | Outdegree | Category |
| --- | --- | --- |
| *Lactobacillus* LMG 17291 | 3289 | STRAIN |
| *Escherichia coli* O157:H7 | 3096 | STRAIN |
| *Pseudomonas aeruginosa* PAO1 | 2074 | STRAIN |
| *Escherichia coli* BL21 | 1474 | STRAIN |
| *Escherichia coli* BL21 DE3 | 1454 | STRAIN |
| *Pseudomonas putida* KT2440 | 1302 | STRAIN |
| *Escherichia coli* MG1655 | 1262 | STRAIN |
| *Escherichia coli* ELI 51 | 1223 | STRAIN |
| *Escherichia coli* K-12 | 1206 | STRAIN |
| *Escherichia coli* Nissle 1917 | 1061 | STRAIN |

**Table 2:** Top 10 nodes by indegree in the phenotypic network.

| Node | Indegree | Category |
| --- | --- | --- |
| LB | 2556 | MEDIUM |
| Biofilm | 2447 | PHENOTYPE |
| Gram negative | 2142 | PHENOTYPE |
| Soil | 2017 | ISOLATE |
| Gram positive | 1981 | PHENOTYPE |
| Human | 1906 | ORGANISM |
| Mouse | 1784 | ORGANISM |
| Probiotic | 1759 | PHENOTYPE |
| LB broth | 1305 | MEDIUM |
| TSB | 1250 | MEDIUM |

The Jaccard index, which computes nodes that are similarly connected to each other within the network, showed mostly closely related strains when looking at the highest values, in some, strain variations for the same species. This showcases the ability of the network to identify relevant properties for entities with similar genetic makeup (Figure 3c). This also works as an internal control for showing the feasibility in identifying reproducible phenotypic traits for different strains. Exceptions to this included *Rhizobium*/*Enterococcus* and *Lactobacillus*/ *Bifidobacterium*, which, although from different phyla, are known to share similar ecological niches i.e., rhizosphere and human gut, respectively (Kumawat et al., 2021; Reuter, 2001).

#### 3.b.ii. Inferring strain-strain correlations from trophic networks

The understanding of microbe-microbe correlations is very relevant for predicting the assemblage and function of microbial communities. However, these correlations are notably hard to study due to experimental constraints, as their inference traditionally entails pairwise growth testing of different strains in the lab and abundance analyses across different samples to study co-occurrence patterns (Freilich et al., 2011; Tackmann et al., 2019). Trophic interactions based on intermediary metabolic compounds have been shown to dictate dynamics in such microbial communities, and can be used as a method for inferring microbe-microbe correlations to complement more classical approaches (Geesink et al., 2024; Gralka et al., 2020).

We used the knowledge network of phenotypes to infer strain-strain correlations by constructing trophic networks based on the LLM predictions from the literature. We built these by grouping strains that inhabited the same hosts or environments according to the INHABITS relation. We next performed an analysis of triads (three nodes contiguously connected) in this subnetwork to count four types of common trophic connections (Figure 4a; see Section 2.b.ii). They included two negative interactions related to competition and inhibition and two positive interactions related to cross-feeding and resistance.

**Figure 4:**
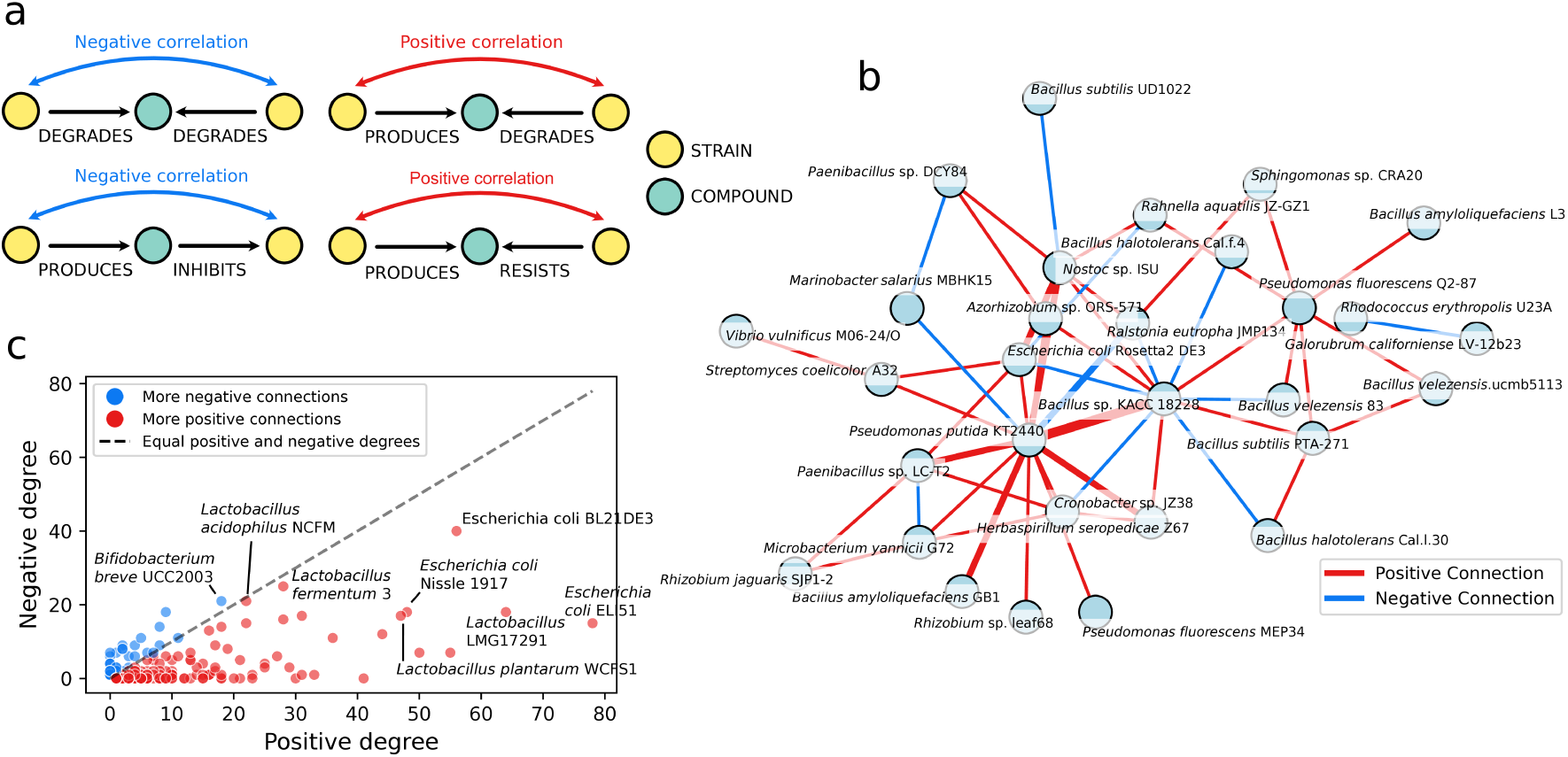
Microbe-microbe correlation inference. **a** Triad combinations in the phenotypic network used to infer trophic microbe-microbe interaction. **b** Inferred strain-strain correlations for bacteria inhabiting *Arabidopsis thaliana* plants. The width of the edges represents the weight of the connection. **c** Node in and out-degree distribution for strain-strain correlations in the inferred trophic interaction network relating to human-colonizing bacteria.

In the constructed strain-strain interaction networks, some nodes were highly connected. For example, when looking at strains associated with *Arabidopsis thaliana* plants, these included strains such as *Pseudomonas putida*, which had many positive correlations both in number and weight (Figure 4b). As a common goods producer, *Pseudomonas putida* has been studied in the context of acting as a driver of microbiome diversity in the rhizosphere of plants (Molina et al., 2019). This shows that these inferred trophic correlations, although biased by the information found in the literature, nevertheless reflect actual biological processes. As found previously, positive connections were more prevalent than negative ones in *A. thaliana*-associated communities, suggesting that community assembly and stability is driven more by cooperation than by competition (Gómez-Pérez et al., 2023).

When looking at other environments such as those associated with humans, which contained a higher number of nodes, we found some strains that were highly interconnected for both positive and negative links to other strains (Figure 4c). Commensal bacteria such as *E. coli* and *Bacillus*, which are generally expected to have a neutral effect towards the host, were the highest when ranking positive correlations. Meanwhile, probiotic bacteria such as *Lactobacillus* and *Bifidobacterium* had a higher ratio of negative to positive connections. This relates to the capacity of probiotics in controlling the microbial community by inhibiting pathogenic competitors (Raheem et al., 2021). Overall, this presents an example of applying the generated phenotypic network to derive biological insights.

### 3.c. Phenotype-gene correlations identify key proteins

#### 3.c.i. Ranking gene importance

In order to find correlations between the genomes of bacteria and their phenotype as predicted by the network, we used gradient boosting. Linking genomes and the phenotype network allowed us to pinpoint specific microbial genes across several taxa as responsible for particular phenotypes. In total, we found ≈9,300 entity-entity significant correlations containing a STRAIN, mainly corresponding to the terms STRAIN-MEDIUM:GROWS_ON (1,774), STRAIN-COMPOUND:PRODUCES (1,573) and COMPOUND-STRAIN:INHIBITS (1,275). This observation partially parallels the relation annotations, further suggesting that increasing the size of the training dataset could lead to a larger number of correlations (Figure 1c). The enriched GO terms for the relevant genes in each of the categories overlapped greatly (Supp. Figure 6). Membrane- and catalytic-activity-related terms were common to most of the relations, highlighting the relevance of proteins that fit these categories in the adaptation and evolution within different environments eventually resulting in different phenotypes (Siliakus et al., 2017; Tomatis et al., 2008). Interestingly, INHIBITS correlations with SPECIES were enriched in biosynthetic terms showing the importance of the production of antimicrobials in such negative correlations (Ochi & Hosaka, 2013).

#### 3.c.ii. Phenotypic traits

The highest importance annotations belonged mostly to PHENOTYPE:PRESENTS categories, but they were varied and did not correlate to the higher abundance entity-entity correlations (Table 3). Additionally, these did not include entity terms that showed the highest degree, indicating that more connections to strains did not necessarily lead to a better significance of correlations. Among the highest importance annotations are known direct correlations between phenotype and functional annotations. For example, many probiotic bacteria are enriched in secreted peptidases including C69 and C1B types, which participate in the breakdown of proteins that improve digestion and feed a healthy gut microbiome (Francavilla et al., 2017). Related to the latter, many probiotics require protein transport machinery to release proteases and other enzymes into the medium, which also contributes to host adhesion and host communication (Sánchez et al., 2008). Accordingly, a protein export membrane protein shows as the second term with highest importance. Other interesting terms that are reported in the literature include PAAR domains in resistance to oxidative stress, in particular for bacteria associated with plants (Li et al., 2024).

**Table 3:** Annotations with importance greater than 150 for their correlation with phenotype. Importance ranking represents the ranking per category of the particular annotation.

| Relation | Term | Gene description | Imp. | Acc. | Rank |
| --- | --- | --- | --- | --- | --- |
| STRAIN-PHENOTYPE:PRESENTS | probiotic | Peptidase C69 | 472.86 | 0.98 | 1 |
| STRAIN-ORGANISM:INHIBITS | *C. albicans* | Peptidase M26, N-terminal domain | 422.29 | 0.96 | 1 |
| STRAIN-MEDIUM:GROWS_ON | PBS-T | Omega transcriptional repressor | 369.31 | 1 | 1 |
| STRAIN-DISEASE:ASSOCIATED_WITH | cystic fibrosis | LysR, substrate-binding | 364.37 | 0.98 | 1 |
| STRAIN-DISEASE:INHIBITS | breast cancer | Uncharacterised protein family, inner membrane CreD | 305.40 | 1 | 1 |
| STRAIN-EFFECT:INHIBITS | binding | Uncharacterised protein family, inner membrane CreD | 290.54 | 1 | 1 |
| STRAIN-PHENOTYPE:PRESENTS | Gram positive | GTP-binding protein OBG, C-terminal | 280.63 | 0.97 | 1 |
| STRAIN-COMPOUND:RESISTS | methicillin | Phenol-soluble modulins beta protein | 269.90 | 0.98 | 1 |
| STRAIN-PHENOTYPE:PRESENTS | probiotic | Protein export membrane protein SecD/SecE, C-terminal | 258.81 | 0.98 | 2 |
| STRAIN-EFFECT:PRESENTS | resistance to oxidative stress | PAAR motif | 225.26 | 1 | 1 |
| STRAIN-SPECIES:INHIBITS | *C. albicans* | LicD/FKTN/FKRP, nucleotidyltransferase domain | 221.03 | 0.96 | 1 |
| STRAIN-COMPOUND:RESISTS | carbapenem | Peptidase M26, N-terminal domain | 219.85 | 0.98 | 1 |
| STRAIN-ORGANISM:INFECTS | mouse | Alkaline shock protein Asp23 | 217.48 | 0.89 | 1 |
| COMPOUND-STRAIN:INHIBITS | tetracycline | Streptococcal surface-anchored protein repeat | 215.89 | 0.99 | 1 |
| STRAIN-EFFECT:INHIBITS | complement activation | Peptidase M26, C-terminal domain | 204.41 | 1 | 1 |
| STRAIN-MEDIUM:GROWS_ON | MRS broth | Exoribonuclease, phosphorolytic domain 1 | 178.69 | 0.99 | 1 |
| STRAIN-ISOLATE:INHABITS | soil | RNA polymerase sigma factor 70, region 4 type 2 | 173.41 | 0.96 | 1 |
| STRAIN-MEDIUM:GROWS_ON | blood agar | Protein of unknown function DUF3290 | 164.57 | 0.99 | 1 |
| STRAIN-PHENOTYPE:PRESENTS | Gram positive | Bacterial surface antigen (D15) | 161.36 | 0.97 | 2 |
| STRAIN-COMPOUND:RESISTS | methicillin | Teichoic acid transporter subunit TagH, SH3-like domain | 157.39 | 0.98 | 2 |
| STRAIN-MEDIUM:GROWS_ON | MRS | Exoribonuclease, phosphorolytic domain 1 | 155.42 | 0.99 | 1 |
| STRAIN-ISOLATE:INHABITS | soil | Glyoxalase/fosfomycin resistance/dioxygenase domain | 154.37 | 0.96 | 2 |
| STRAIN-COMPOUND:RESISTS | carbapenem | Periplasmic binding protein/LacI sugar binding domain | 151.36 | 0.98 | 2 |

In some cases, we found the same highest importance annotation for two different categories with the same meaning, e.g., presents a cellulolytic phenotype and degrades metabolite cellulose (Table 4), which resulted in similar Glycoside Hydrolase (GH) annotations. GH9 family proteins in particular include enzymes such as endoglucanase and cellobiohydrolases, which are known for their cellulose degradation activities (Konar et al., 2022; Liu et al., 2011). The ability of different models to converge in the same correlation shows the robustness of the method and the ability of the approach to retrieve biologically-sound information related to phenotypic traits.

**Table 4:** High-importance cellulose-related annotations in the gene-phenotype correlation dataset.

| Gene description | Term | Relation | Imp. | Rank |
| --- | --- | --- | --- | --- |
| cellulose | Cellulose-complementing protein | STRAIN-COMPOUND:PRODUCES | 64.80 | 1 |
| cellulose | Glycoside hydrolase family 9 | STRAIN-COMPOUND:DEGRADES | 40.17 | 1 |
| cellulose | Protein of unknown function DUF2501 | STRAIN-COMPOUND:PRODUCES | 28.14 | 2 |
| cellulose | Carbohydrate binding module family 6 | STRAIN-COMPOUND:DEGRADES | 27.26 | 2 |
| cellulose | Cytochrome c-type biogenesis protein CcmB | STRAIN-COMPOUND:PRODUCES | 22.14 | 3 |
| cellulolytic | Glycoside hydrolase family 9 | STRAIN-PHENOTYPE:PRESENTS | 19.17 | 1 |
| cellulose | Haem-binding domain | STRAIN-COMPOUND:PRODUCES | 18.88 | 4 |
| cellulose | Aminopeptidase N-like, N-terminal domain | STRAIN-COMPOUND:DEGRADES | 17.99 | 3 |
| cellulolytic | Glycoside hydrolase family 10 domain | STRAIN-PHENOTYPE:PRESENTS | 16.11 | 2 |
| cellulose | Inner membrane-spanning protein YciB | STRAIN-COMPOUND:DEGRADES | 15.11 | 4 |

#### 3.c.iii. Host-association and pathogenicity

To effectively study genes associated with host colonization and infection, we examined these in our correlation study. Our approach allows us to identify genes that may be relevant for adapting to specific hosts. For the term STRAIN-ORGANISM:INHABITS, we found a significant phylogenetically diverse set of organism hosts, including animals and plants (Supp. Figure 6). The lack of overlap in high-importance genes among hosts likely relates to the different strategies that bacteria use to adapt to a particular host, and also to the large phylogenetic diversity of hosts in the dataset. However, we found some in common for related organisms, these included an RNA polymerase sigma factor for two crops, rice and tomato, as well as CsgD-like domain for shrimp and algae, which share similar environmental conditions. The former may be related to stress response (Paget & Helmann, 2003) and the latter to biofilm formation, which may be relevant in the colonization of both hosts (Gerstel & Römling, 2003; Ogasawara et al., 2011).

Regarding the plant-host high importance genes, we found many that are known to be relevant for symbiotic relationships. These included glutathione transferase in soybean, which is involved in adapting to biotic stresses associated with plant-colonization (Gullner et al., 2018) and AvrE effector domains for potato which interact with the plant and help dampen the defense response to promote succesful settling on the root (Degrave et al., 2015).

Performing a similar approach on the ORGANISM:INFECTS relation, we also noticed a lack of overlap. However, when looking at the annotations, we found more terms directly relating to pathogenicity and not necessarily symbiotic or neutral effect on the host, suggesting that the models are able to capture this difference. Mice were one of the organisms with the larger number of high importance associations, highlighting the use of these as animal models in disease research (Supp. Figure 7). One of the microbial genes related to infection in mice with the highest importance was OmpA-like domain, which is a known virulence factor that has been studied in *E. coli* (Selvaraj et al., 2007). Two of the three top genes, namely OmpA-like domains and GHs, appear as highly significant in BacFITBase, created by experimentally measuring pathogenic persistence in mice by several model pathogens (Rendón et al., 2019). Similarly, the top results for other hosts, including chicken (MaoC-like dehydratase) and rabbit (short-chain dehydrogenase/reductase), also show highly significant matches.

Other potential virulence factors that have been reported in literature present in this dataset include pullulanase (Uda et al., 2016), surface antigen (Blanco et al., 1997), Shiga-like toxin (Beutin et al., 1995), histidine kinases (Bem et al., 2015). Given the prevalence of these, we hypothesize that the other enzymes that do not directly fit the virulence paradigm may present potential virulence factors indirectly advancing pathogenicity in a particular host. These included globin-sensor domain and anti-bacteriophage protein A/HamA, with known roles in environmental sensing and defense against bacteriophages (Stranzl et al., 2011; Yasui et al., 2014), respectively.

Antimicrobial compound production and defense also feature heavily in the subset of high-importance genes related to host colonization in both categories. This included production of antimicrobial peptides such as lantibiotic (Riley & Wertz, 2002), bacteriocin (Gabrielsen et al., 2014), sublancin (Wu et al., 2019) and colicin (Chatterjee et al., 2005). More indirect methods of suppression were also found such as beta-lactamase inhibitor production (Docquier & Mangani, 2018). Bacterial biocontrol compounds are important for settling hosts as they free up currently occupied niches and prevent competition by other bacteria (García-Bayona & Comstock, 2018). Regarding antimicrobial resistance, there are terms including tetracycline repressor (Orth et al., 2000), glyoxalase/fosfomycin resistance (Shen et al., 2010), histidine kinase (Bem et al., 2015), outer membrane protein transport protein (OMPP1/FadL/TodX) (Kesavan et al., 2020), ABC transporters (Greene et al., 2018) and efflux pumps (Poole, 2007). The prevalence of terms belonging to this category in both host-associated categories highlights the current problem of antibiotic resistance development in fighting infection. Interesting is also the abundance of proteins with unknown function which suggest potential novel connections, as well as demonstrate the reliance of this method on accurate functional annotations. Nonetheless, uncharacterized genes regarding their function present potential candidates for a more targeted study.

#### 3.c.iv. Positive selection in highly important features

When looking at a subset of these correlations and their highest importance genes, we saw an enrichment in genes under positive selective pressures. For all the genes tested, 2,630 (30.7%) showed evidence for positive selection. This is much higher than the predicted base rate for genes under detectable selection in a particular genome, which is variable but does not generally exceed 20% (Malekian et al., 2022; Richards et al., 2019). The prevalence of positive selection underscores the relevance of these highly correlated genes to particular phenotypes in adaptation. Overall, there was a small positive correlation between the phenotype importance values and the evidence for positive selection as measured with the likelihood ratio test, and the top genes belonged mainly to the STRAIN-COMPOUND:PRODUCES and STRAIN-PHENOTYPE:PRESENTS categories (Figure 5a, Table 5).

**Figure 5:**
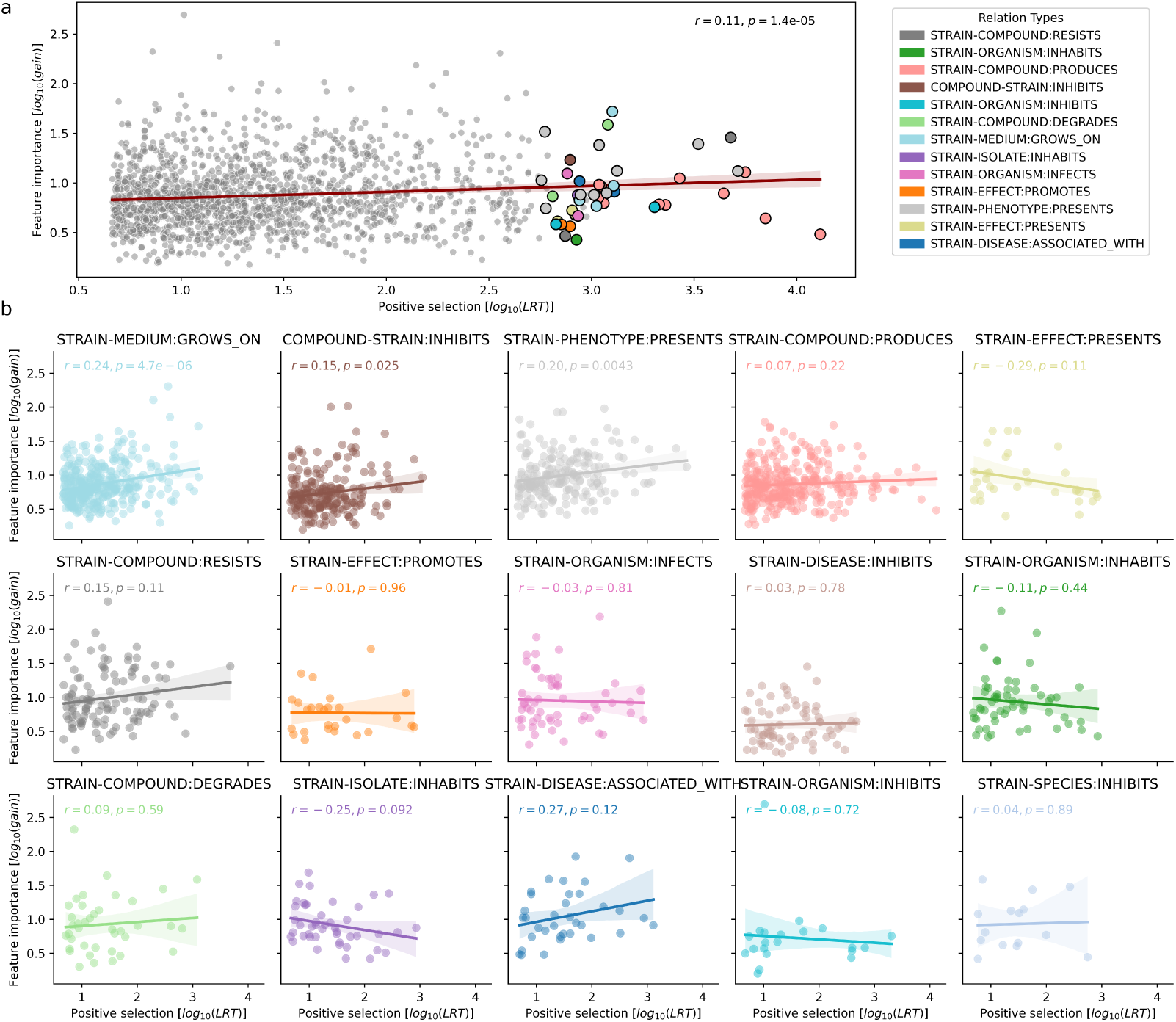
Positive selection in relevant genes. **a** Positive pressure correlation among the top importance annotations and the likelihood ratio test (LRT) for all significant protein families with evidence of positive selection. Correlations where log10(LRT) > 2.75 are highlighted with a color representing their relation type. **b** Positive pressure correlation among the top importance annotations and the likelihood ratio test (LRT) for positive selection of the highest populated relation categories containing at least 30 gene correlations.

**Table 5:** Gene-phenotype correlations with high evidence of positive selection. The logLRT and logImp. columns represent the log-likelihood ratio test and the log importance values, respectively. Top 3 genes where logLRT > 2.5 for each category are depicted.

| Gene description | Term | Relation | logLRT | logImp. |
| --- | --- | --- | --- | --- |
| Domain of unknown function DUF6708 | APE | STRAIN-COMPOUND:PRODUCES | 4.11 | 0.48 |
| STAS domain | hopene | STRAIN-COMPOUND:PRODUCES | 3.85 | 0.64 |
| PRC-barrel domain | alkaliphilic | STRAIN-PHENOTYPE:PRESENTS | 3.71 | 1.12 |
| Putative lipoprotein, pathogenic bacteria | rifampicin | STRAIN-COMPOUND:RESISTS | 3.68 | 1.46 |
| Glycoside hydrolase family 9 | cellulolytic | STRAIN-PHENOTYPE:PRESENTS | 3.52 | 1.39 |
| Gamma-butyrobetaine hydroxylase-like, N-terminal | *Rhizoctonia solani* | STRAIN-ORGANISM:INHIBITS | 3.31 | 0.76 |
| Mub B2-like domain | candidiasis | STRAIN-DISEASE:ASSOCIATED_WITH | 3.11 | 0.91 |
| Aminotransferase, class I/classII, large domain | BA | STRAIN-MEDIUM:GROWS_ON | 3.11 | 0.97 |
| Phenol-soluble modulin beta protein | TSB | STRAIN-MEDIUM:GROWS_ON | 3.10 | 1.72 |
| CBS domain | cellulose | STRAIN-COMPOUND:DEGRADES | 3.08 | 1.59 |
| Serine hydroxymethyltransferase-like domain | BV | STRAIN-DISEASE:ASSOCIATED_WITH | 2.94 | 1.02 |
| Sarcosine oxidase, delta subunit, heterotetrameric | *Arabidopsis* | STRAIN-ORGANISM:INFECTS | 2.93 | 0.67 |
| GGDEF domain | carious dentine | STRAIN-ISOLATE:INHABITS | 2.93 | 0.88 |
| Protein of unknown function DUF333 | ginger | STRAIN-ORGANISM:INHABITS | 2.93 | 0.43 |
| C883_1060-like, ketoreductase domain, N-terminal | rhizosphere colonization | STRAIN-EFFECT:PRESENTS | 2.92 | 0.68 |
| Stage II sporulation protein M-like | antifungal action | STRAIN-EFFECT:PRESENTS | 2.91 | 0.72 |
| KfrB domain | ceftriaxone | COMPOUND-STRAIN:INHIBITS | 2.90 | 1.23 |
| Protein of unknown function DUF6060 | *Nicotiana benthamiana* | STRAIN-ORGANISM:INFECTS | 2.88 | 1.10 |
| Methionyl/Valyl/Leucyl/Isoleucyl-tRNA synthetase, anticodon-binding | FA | STRAIN-COMPOUND:RESISTS | 2.87 | 0.47 |
| Glutamate/phenylalanine/leucine/valine/L-tryptophan dehydrogenase, C-terminal | drought stress tolerance | STRAIN-EFFECT:PROMOTES | 2.85 | 0.58 |
| Beta-lactamase-related | *Fusarium oxysporum* | STRAIN-ORGANISM:INHIBITS | 2.83 | 0.58 |
| Pyruvate formate lyase domain | lignin | STRAIN-COMPOUND:DEGRADES | 2.81 | 0.87 |
| Apiosidase-like, catalytic domain | *Serratia marcescens* | STRAIN-SPECIES:INHIBITS | 2.74 | 0.44 |
| Glycosyl hydrolase family 38, C-terminal | depression | STRAIN-DISEASE:INHIBITS | 2.67 | 0.64 |
| Polyketide synthase, dehydratase domain, N-terminal | *L. minor* | STRAIN-ORGANISM:INHABITS | 2.64 | 0.52 |
| Sodium-dependent phosphate transport protein | feces | STRAIN-ISOLATE:INHABITS | 2.60 | 0.81 |
| Domain of unknown function (DUF6895) | BPH | STRAIN-DISEASE:INHIBITS | 2.58 | 0.65 |

Examples of genes under strong positive selection in this dataset that were also highly correlated to phenotypes included GH9, which was linked to a cellulolytic phenotype. Cellulose degradation is relevant in many contexts of bacterial survival including fiber breakdown in the gut of organisms that feed on plant material (Ali et al., 2019; Weimer, 1992) and other mutualistic microbe-microbe interactions in soil (Soundar & Chandra, 1987). Another type of protein domain in the top genes is beta-lactamase-related, which has been also found under high selective pressure (Gniadkowski, 2008) and is involved in antibiotic resistance. This is expected as antibiotics are a strong pressure behind much of the adaptive selection related to competition mechanisms, particularly against *Fusarium oxysporum*, which is also known to produce beta-lactamases (Chang et al., 2021). We grouped these correlations and found that for the most populated categories there was a significant positive correlation to highest importance genes, including STRAIN-MEDIUM:GROWS_ON, COMPOUND-STRAIN:PRESENTS and STRAIN-PHENOTYPE:PRESENTS. This suggests that diversifying selection is driving adaptation behind particular phenotypes including adaptation to media and inhibition. In the other categories, although mostly a positive trend, there was no significant pattern of correlation (Figure 5b). This lack of significant positive correlation could be explained by the bias in prediction and the differences in the distribution of RE catogories tested for positive selection.

## 4. Conclusion

This work demonstrates NLP’s potential to unlock rich bacterial phenotype information from scientific literature. The constructed phenotype network revealed a power-law distribution with hubs consisting of well-studied strains and common phenotypes, highlighting the ability of the network to identify reproducible traits as well as potential biases in research focus. This emphasizes the importance of more research on less studied and exotic strains, allowing future predictions to be more generalizable. Nonetheless, the network enabled trophic network inference from strains associated with various environments, in order to assess microbe-microbe interactions. The gradient boosting analysis successfully inferred phenotype-gene correlations, agreeing with the current scientific literature and uncovering novel patterns related to traits, pathogenicity, and host-association. Notably, we found an enrichment in antimicrobial production and defense proteins for host-association and pathogenicity. Many of these novel findings could be useful for experimental researchers as they allow potential candidate genes to be identified when investigating certain phenotypes, presenting a valuable resource for the scientific community. Importantly, instances of positive selection highlighted genes underlying adaptive phenotypes in two of the relation categories.

Comparing our results with the bacterial phenotype database BacDive (Söhngen et al., 2014), we find a lower number of strains with annotated characteristics (≈60,000 vs. ≈97,000 as of current release). However, we find these more evenly taxonomically spread in our dataset, compared to the database which is heavily biased towards *Streptomyces* (1/4 of total strains). While our approach is less reliable to create accurate annotations and more sensitive to publication bias than manual curation, it has the potential to scale better and become more complete as the corpus grows and computational capacity improves. Additionally, our approach is restricted to qualitative data and so lacks the nuances of quantitative measures, such as optimal growth temperature or pH. These may be viewed as benefits since they allow experimental researchers to easily obtain a snapshot of the phenotype information of their strain of interest as it appears in the literary corpus. This can be very advantageous for sensibly directing research resources to solve specific problems. Moving forward, improvements could be made to the network analysis by taking into account taxonomy of the strains and confidence weight by independently reported traits.

In summary, this work paves the way for large-scale, automated exploration of the relationships between microbial genotype and phenotype, and provides some examples and strategies to analyze such extensive data.

## Acronyms

NLP: Natural Language Processing
LLMs: Large Language Models
RE: Relation Extraction
NER: Named Entity Recognition
PMC: PubMed Central
BERT: Bidirectional Encoder Representations from Transformers

---

**Supp. Figure 1:**
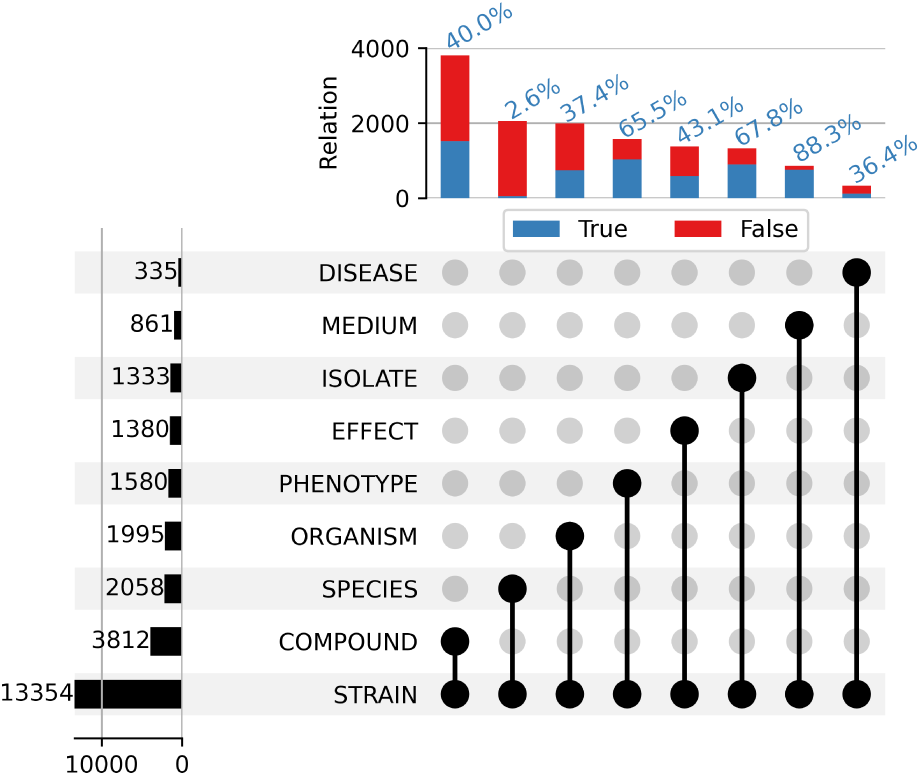
Upset group plot showing the sentence co-occurrence of STRAIN labels with others in the annotated dataset. Labeled in blue on top of the bars is the percentage of entities with a positive presence of a relation extraction (RE) annotation among them.

**Supp. Figure 2:**
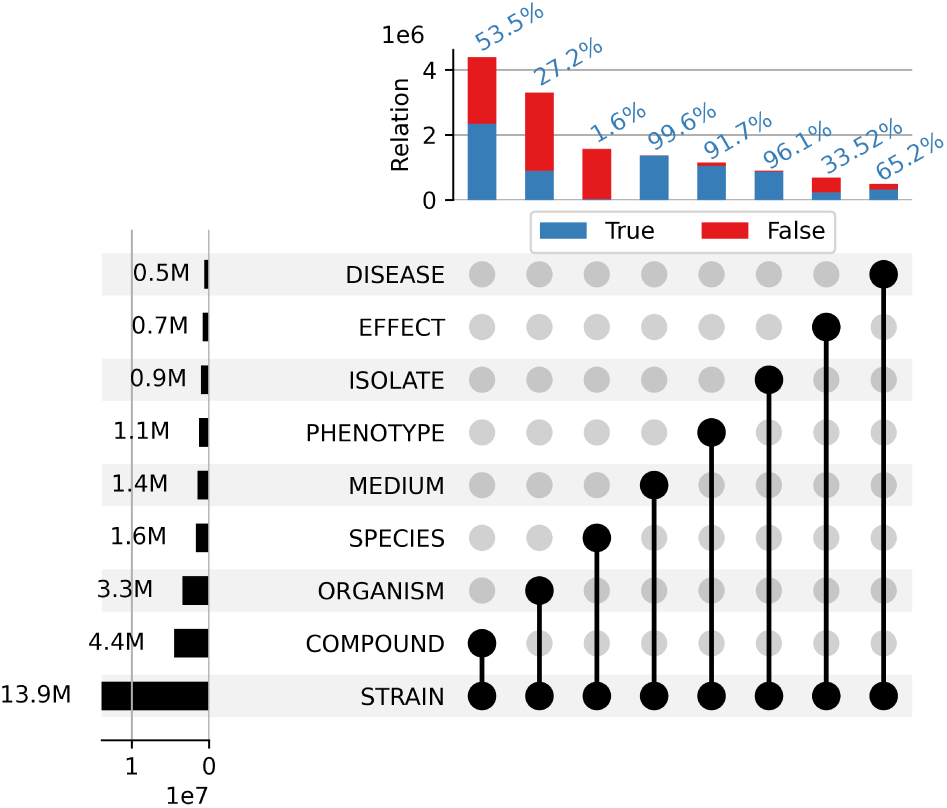
Overview of predictions in the PMC corpus. Upset group plot showing the sentence co-occurrence of STRAIN labels with others in the predicted dataset. Labeled in blue on top of the bars is the percentage of entities with a positive presence of a relation extraction (RE) annotation among them.

**Supp. Figure 3:**
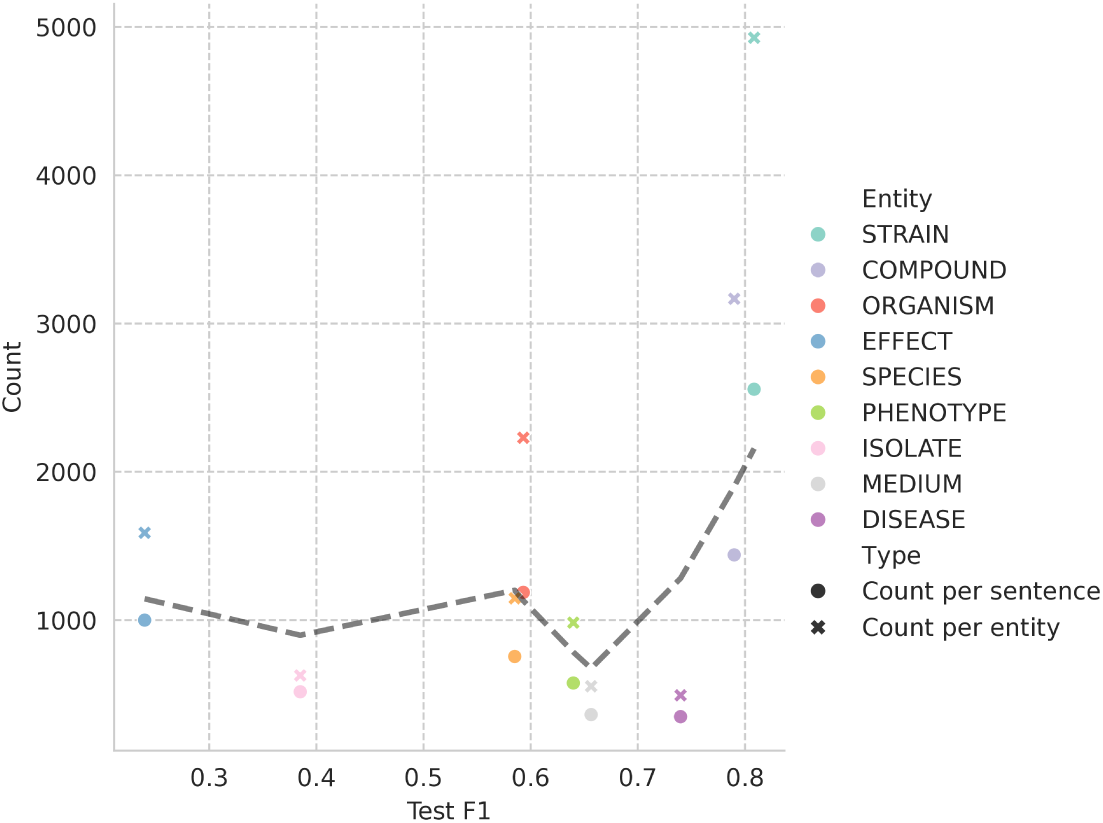
Correlation of the number of labeled entities in the annotation dataset to model performance. The counts by either sentence, circle, or entity, cross, are displayed correlated to the F1 performance on the test set after training grouped by the colored entity. Trend line is drawn using LOESS smoothing over all points.

**Supp. Figure 4:**
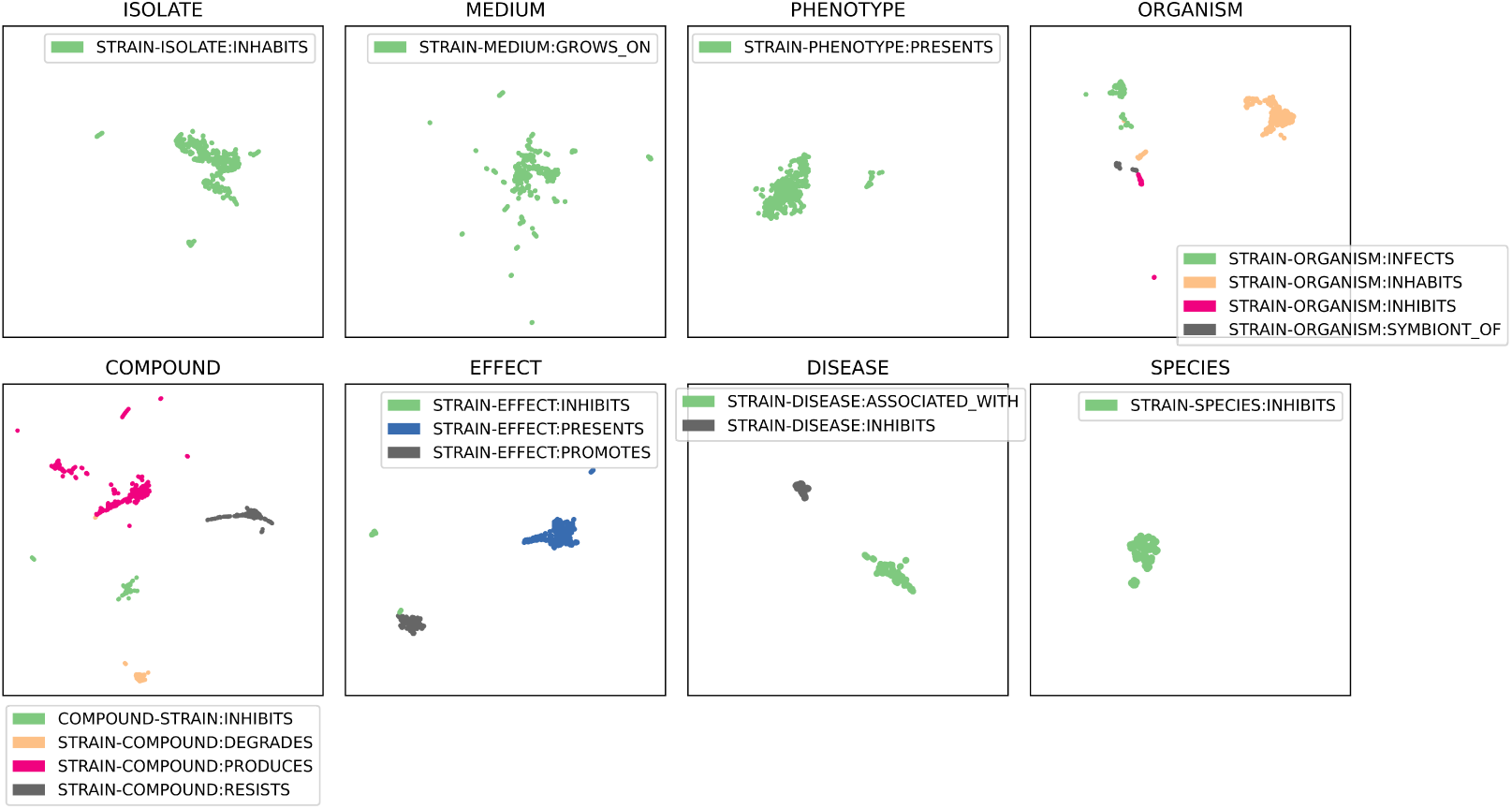
UMAP representation of sentence embeddings in the training set for each of the different relation extraction models grouped by entity. Displayed are only sentences that are positive for the presence of that particular relation. Axes correspond to UMAP dimensions 1 and 2.

**Supp. Figure 5:**
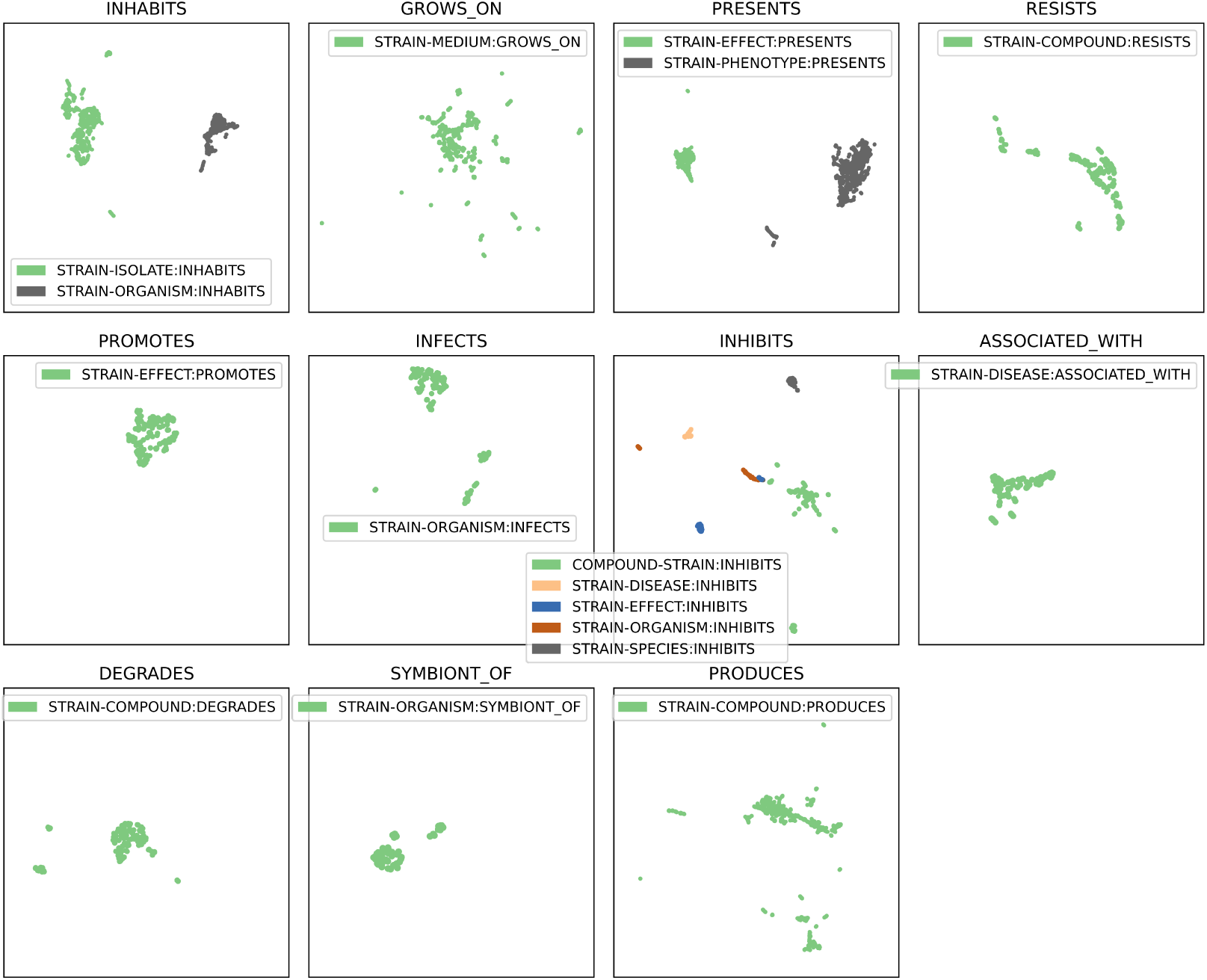
UMAP representation of sentence embeddings in the training set for each of the different relation extraction models grouped by relation. Displayed are only sentences that are positive for the presence of that particular relation. Axes correspond to UMAP dimensions 1 and 2.

**Supp. Figure 6:**
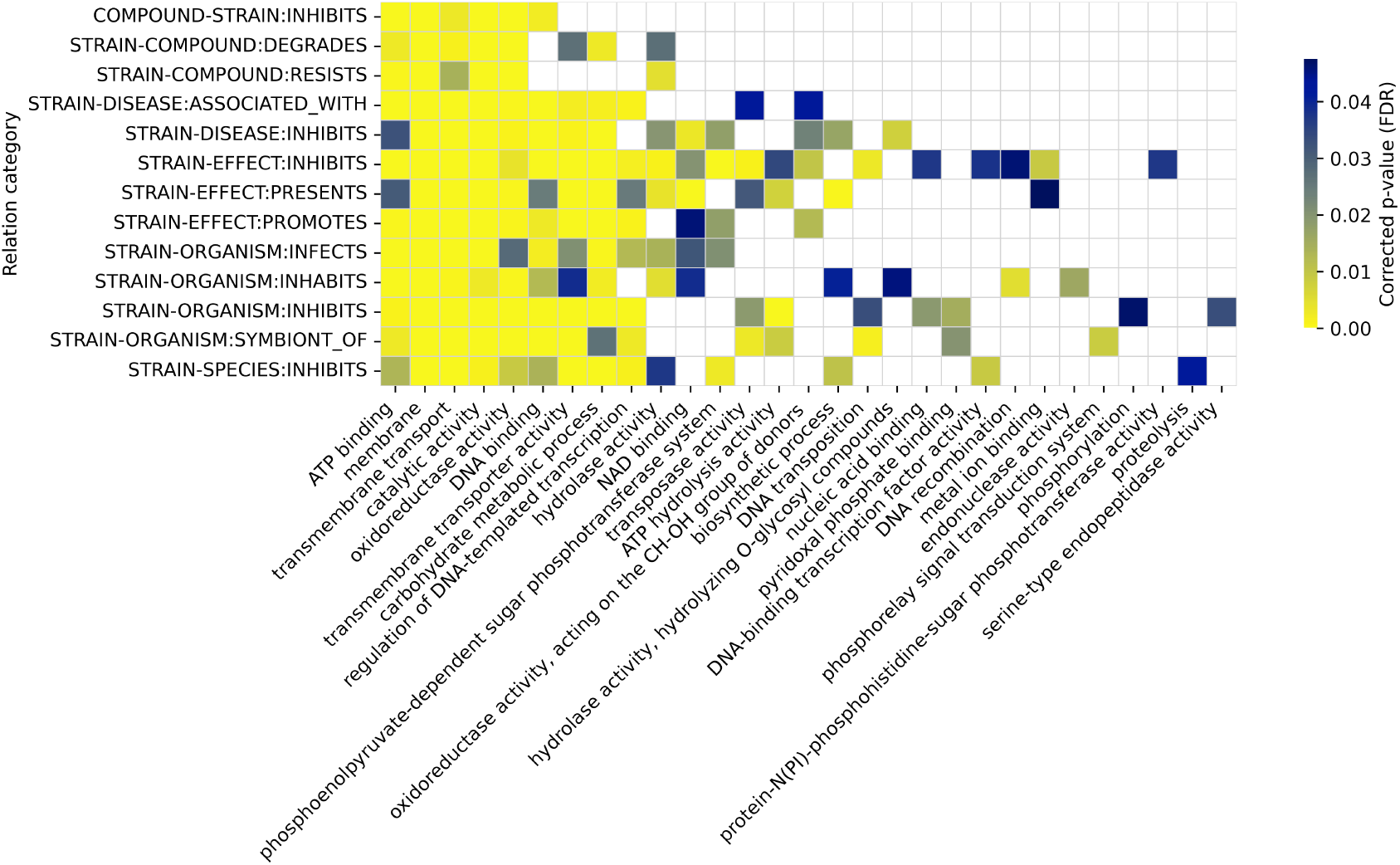
GO enrichment of high-importance genes in the correlation to phenotype and grouped by their relation category.

**Supp. Figure 7:**
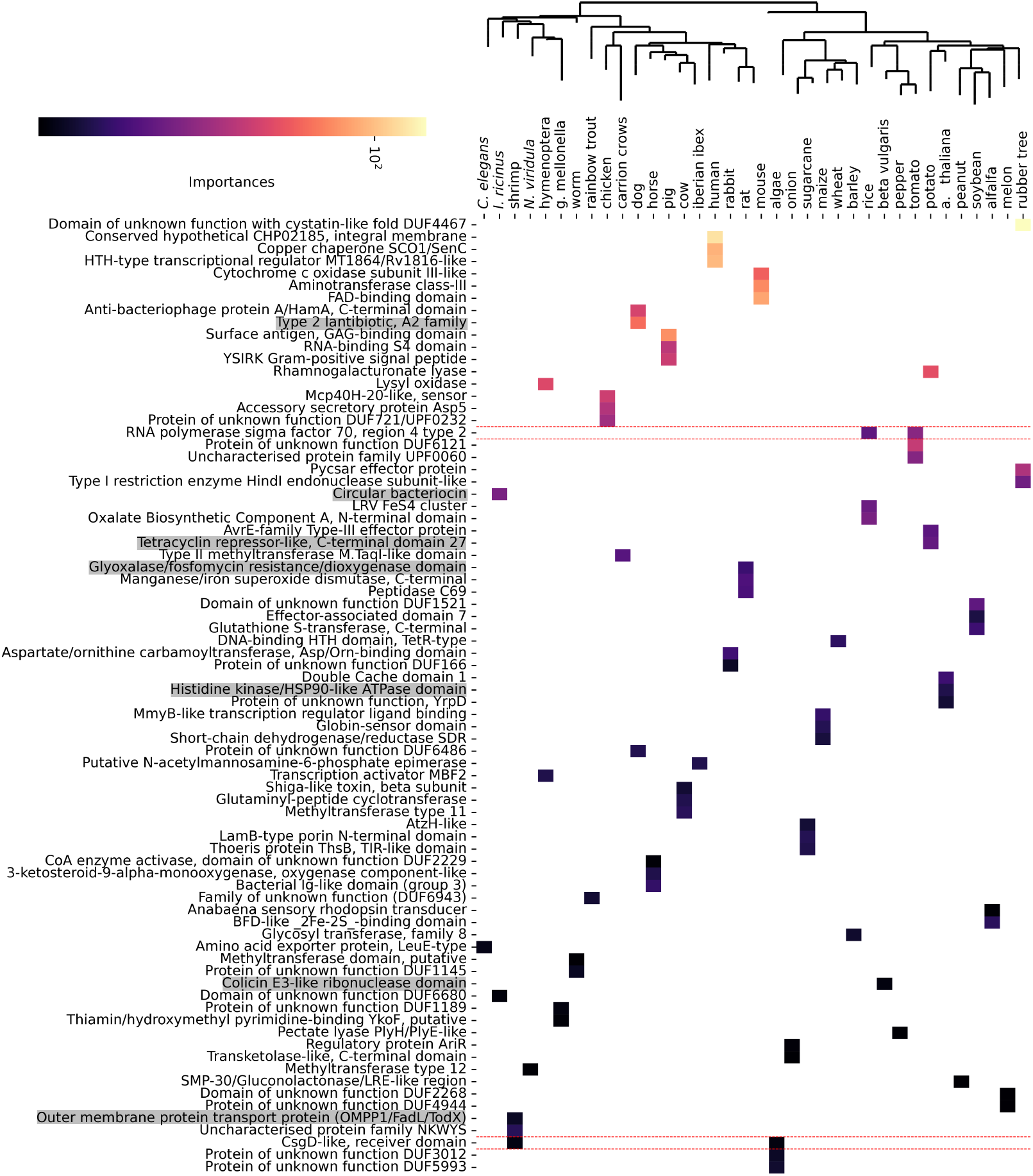
Prevalence of bacterial genes in the STRAIN-ORGANISM:INHABITS ORGANISM relations. At the top, the phylogenetic tree represents the distance between the hosts. Highlighted with red lines are the terms that occur in more than one host. Highlighted genes in grey correspond to antimicrobial production or resistance.

**Supp. Figure 8:**
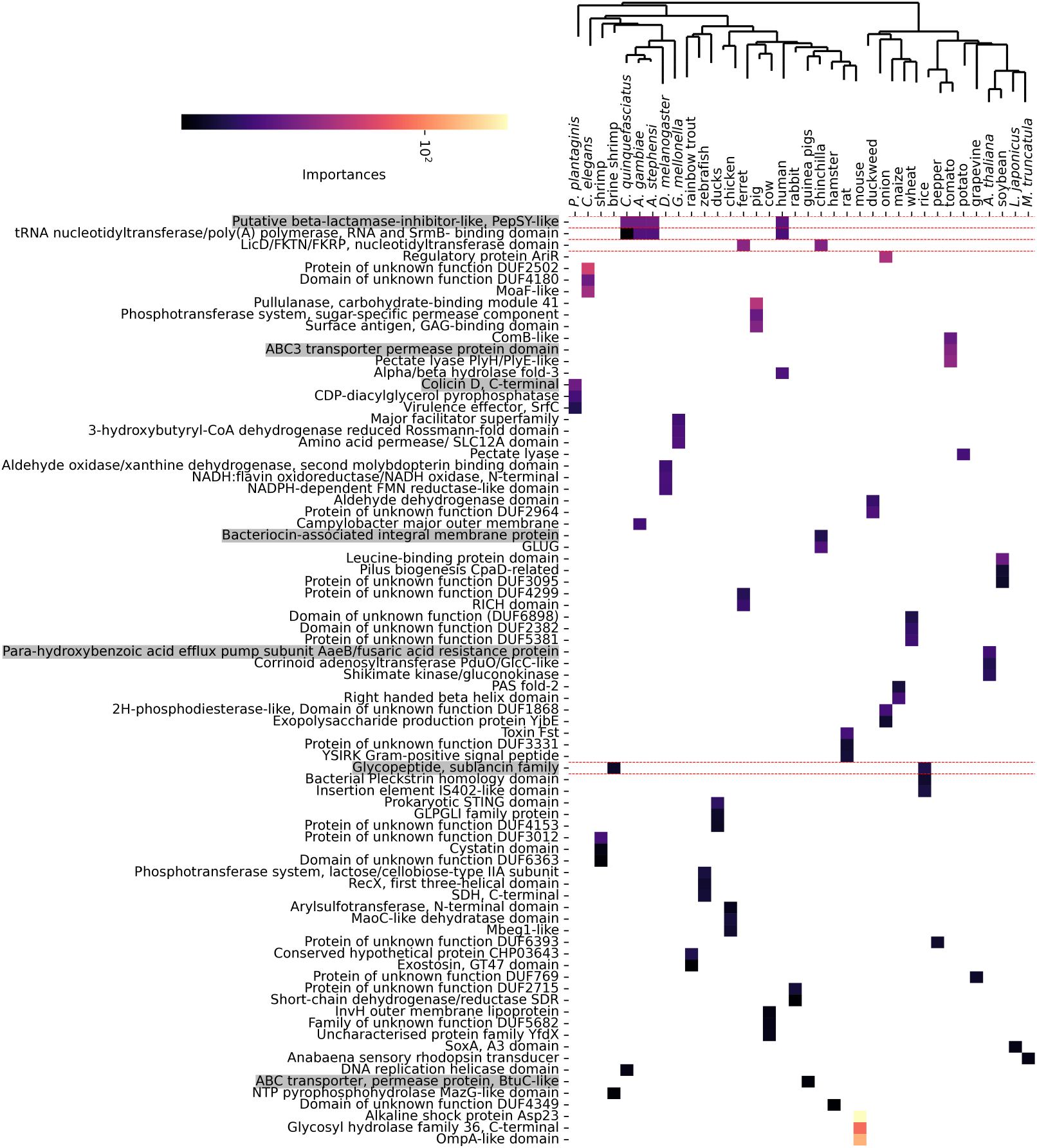
Prevalence of bacterial genes in the STRAIN-ORGANISM:INFECTS relations. At the top, the phylogenetic tree represents the distance between the hosts. Highlighted with red lines are the terms that occur in more than one host. Highlighted genes in grey correspond to antimicrobial production or resistance.

